# High dimensional analyses of cells dissociated from cryopreserved synovial tissue

**DOI:** 10.1101/284844

**Authors:** Laura T. Donlin, Deepak A. Rao, Kevin Wei, Kamil Slowikowski, Mandy J. McGeachy, Jason D. Turner, Nida Meednu, Fumitaka Mizoguchi, Maria Gutierrez-Arcelus, David J. Lieb, Joshua Keegan, Kaylin Muskat, Joshua Hillman, Cristina Rozo, Edd Ricker, Thomas M. Eisenhaure, Shuqiang Li, Edward P. Browne, Adam Chicoine, Danielle Sutherby, Akiko Noma, Accelerating Medicines Partnership: RA/SLE Network, Chad Nusbaum, Stephen Kelly, Alessandra B. Pernis, Lionel B. Ivashkiv, Susan M. Goodman, William H. Robinson, Paul J. Utz, James A. Lederer, Ellen M. Gravallese, Brendan F. Boyce, Nir Hacohen, Costantino Pitzalis, Peter K. Gregersen, Gary S. Firestein, Soumya Raychaudhuri, Larry W. Moreland, V. Michael Holers, Vivian P. Bykerk, Andrew Filer, David L. Boyle, Michael B. Brenner, Jennifer H. Anolik

## Abstract

**Background:** Detailed molecular analyses of cells from rheumatoid arthritis (RA) synovium hold promise in identifying cellular phenotypes that drive tissue pathology and joint damage. The Accelerating Medicines Partnership (AMP) RA/SLE network aims to deconstruct autoimmune pathology by examining cells within target tissues through multiple high-dimensional assays. Robust standardized protocols need to be developed before cellular phenotypes at a single cell level can be effectively compared across patient samples.

**Methods:** Multiple clinical sites collected cryopreserved synovial tissue fragments from arthroplasty and synovial biopsy in a 10%-DMSO solution. Mechanical and enzymatic dissociation parameters were optimized for viable cell extraction and surface protein preservation for cell sorting and mass cytometry, as well as for reproducibility in RNA sequencing (RNA-seq). Cryopreserved synovial samples were collectively analyzed at a central processing site by a custom-designed and validated 35-marker mass cytometry panel. In parallel, each sample was flow sorted into fibroblast, T cell, B cell, and macrophage suspensions for bulk population RNA-seq and plate-based single cell CEL-Seq2 RNA-seq.

**Results:** Upon dissociation, cryopreserved synovial tissue fragments yielded a high frequency of viable cells, comparable to samples undergoing immediate processing. Optimization of synovial tissue dissociation across six clinical collection sites with ∼30 arthroplasty and ∼20 biopsy samples yielded a consensus digestion protocol using 100µg/mL of Liberase TL^™^ enzyme. This protocol yielded immune and stromal cell lineages with preserved surface markers and minimized variability across replicate RNA-seq transcriptomes. Mass cytometry analysis of cells from cryopreserved synovium distinguished: 1) diverse fibroblast phenotypes, 2) distinct populations of memory B cells and antibody-secreting cells, and 3) multiple CD4+ and CD8+ T cell activation states. Bulk RNA sequencing of sorted cell populations demonstrated robust separation of synovial lymphocytes, fibroblasts, and macrophages. Single cell RNA-seq produced transcriptomes of over 1000 genes/cell, including transcripts encoding characteristic lineage markers identified.

**Conclusion:** We have established a robust protocol to acquire viable cells from cryopreserved synovial tissue with intact transcriptomes and cell surface phenotypes. A centralized pipeline to generate multiple high-dimensional analyses of synovial tissue samples collected across a collaborative network was developed. Integrated analysis of such datasets from large patient cohorts may help define molecular heterogeneity within RA pathology and identify new therapeutic targets and biomarkers.

## Background

The destructive inflammatory environment in rheumatoid arthritis (RA) synovium results from the activity of various cell types, including synovial fibroblasts, macrophages, lymphocytes, osteoclasts, and vascular endothelial cells [1–5]. Multiple pathways can be targeted to treat rheumatoid arthritis, including inhibition of tumor necrosis factor (TNF) or interleukin-6 (IL-6) signaling, blockade of T cell costimulation, depletion of B cells, and inhibition of the JAK/STAT pathway [6]. However, despite the advent of biologic therapies, up to ∼2/3 of RA patients do not achieve sustained disease remission [7], and there are no reliable biomarkers that serve to guide selection of specific therapeutic options for the individual patient. A more comprehensive interrogation of cells present in rheumatoid synovium may identify additional pathologic pathways targetable by therapeutics and aid in stratifying patients into disease subsets and treatment response categories based on informative biomarkers [8, 9]. Such studies have previously been limited by the lack of methods to simultaneously recover and assay the diversity of cell types in the synovium and by the challenge of applying high dimensional single cell analytics to sufficient numbers of patient samples. In addition, comparisons of identical cell lineages isolated from targeted tissues in patients with RA and other diseases such as systemic lupus erythematosus (SLE) have not been performed.

In recognition of these needs, the NIH Accelerating Medicines Partnership (AMP) RA/SLE network was assembled with the goal of generating detailed analyses of synovial tissue samples from a large number of RA patients. The RA Working Group of the AMP RA/SLE Network (or AMP RA network) has developed a pipeline to study RA synovial tissue samples through parallel high-dimensional analyses including mass cytometry, bulk RNA-seq of selected cell populations, and single cell RNA-seq. Mass cytometry can define the cellular landscape of tissues, while RNA sequencing delves deeper into the gene expression profile for each cell type. When performed at scale on a large patient cohort, these assays have the potential to aid in developing personalized therapeutic approaches for RA.

To amass an RA cohort of sufficient size, numerous clinical collection sites contributed synovial tissue samples. The network set out to develop a pipeline for uniform and reproducible sample handling involving cryopreservation of intact synovial tissue samples. The frozen tissue fragments were then shipped to a central processing site capable of performing multiple high-dimensional analyses in parallel for each sample. The protocol was optimized for two sources of tissue: synovial biopsy obtained for research and clinically-indicated excision during arthroplasty.

Here, we report the AMP RA network protocol for dissociation and analysis of immune and stromal cells from cryopreserved synovial tissue by multiple high-dimensional technologies. We describe a consensus protocol for synovial tissue disaggregation that results in high yields of viable cells with preserved surface marker expression—for both larger arthroplasty and millimeter sized biopsy samples. We also describe an experimental pipeline to analyze each sample in parallel using single-cell RNA-seq and low-input bulk RNA-seq on selected populations, as well as mass cytometry with defined markers for synovial cells. This pipeline allows for cytometric identification and quantification of numerous immune and stromal cell populations, as well as robust transcriptomic analyses of small populations of cells from pathologic synovial samples. These methods are adaptable for use in multiple sites and have been adopted and utilized by the AMP RA network.

## Methods

### Human patient samples

Patients with RA fulfilled the ACR 2010 Rheumatoid Arthritis classification criteria or were identified by the participating site rheumatologist as having clinical RA [6]. A diagnosis of osteoarthritis (OA) was defined by the treating surgeon and confirmed by chart review. Synovial tissue samples were acquired by two different methods: arthroplasty and ultrasound-guided synovial biopsy. Arthroplasty samples were acquired after removal as part of standard of care at four U.S. institutions (Hospital for Special Surgery, NY, University California San Diego, CA, University of Pittsburgh, PA, and University of Rochester, NY). All synovial biopsy samples were acquired from clinically inflamed joints under research protocols at the University of Birmingham, UK and Barts & the London School of Medicine and Dentistry, UK. The study received institutional review board approval at each site.

### Arthroplasty synovial tissue collection

Synovial tissue excised as standard of care during arthroplasty was transported from the operating room to a laboratory in ice-cold PBS. The tissue was identified as synovium by characteristic features: a fibrous or elastic tissue, pink or reddish in appearance. Small fragments (∼1-2mm^3^) were generated by dissecting with forceps or surgical scissors. Six fragments (∼150mg total) across the tissue were randomly combined to make individual aliquots [10, 11].

### Synovial biopsy procedures

Synovial biopsy samples were obtained under ultrasound guidance by an experienced interventionist using either a needle biopsy or portal-and-forceps. For needle biopsy procedures, a 14G or 16G spring loaded Cook Quick-Core® Biopsy needle was used to obtain multiple fragments [12, 13]. For the portal-based approach, a portal (Merit Medical Prelude 7F) was initially inserted into the synovial cavity with multiple samples then retrieved using flexible 2.0-2.2mm forceps [14, 15]. Six biopsy fragments were randomly allocated per aliquot.

### Synovial tissue cryopreservation and thawing

Synovial tissues were either (1) disaggregated immediately, followed by immediate cryopreservation of dissociated cells; with the dissociated cells cryopreserved or (2) cut into fragments that were cryopreserved for subsequent disaggregation at a central processing site. Dissociated synovial cells were resuspended in CryoStor® CS10 (BioLife Solutions) at ∼2 million cells/mL to viably freeze them. Intact synovial tissue samples were divided into fragments as described above and transferred to a cryovial (1.5mL, Nalgene) containing 1mL of CryoStor® CS10 for viable freezing. Cryovials were then placed into an insulated container with isopropanol in the bottom chamber for slow freezing (Mr. Frosty, Nalgene), which comprised incubation at 4°C for 10 minutes followed by one day at −80°C. The samples were then either shipped on dry ice or transferred into liquid nitrogen for long-term storage.

Synovial fragments were thawed by rapidly warming the cryovial in a 37°C water bath. The preservation media was filtered out through a 70µm strainer. The tissue was then rinsed through a series of incubations in a 6-well culture plate; 10 minutes in 10% FBS/RPMI at room temperature with intermittent swirling, a quick rinse in 10% FBS/RPMI and a final rinse in serum-free RPMI. Frozen synovial cells were thawed rapidly in a 37°C water bath and transferred into 20mL of 10% FBS/RPMI, centrifuged to pellet cells, and then resuspended in media for downstream analyses.

### Dissociation of synovial tissue

Both arthroplasty and synovial biopsy samples were dissociated by a combination of enzymatic digestion and mechanical disruption with various test conditions described in the Results section. The final consensus AMP RA network enzyme digestion uses RPMI media with Liberase TL^™^ enzyme preparations (100µg/mL, Roche) and DNaseI (100µg/mL, Roche).

#### Arthroplasty tissue

To achieve consistent mechanical disruption and proteolytic enzyme exposure across a fragment of tissue, large synovial specimens (>150mg—gram) were cut into small fragments (∼2mm^3^) using surgical scissors. Mechanical disruption of arthroplasty samples (six tissue fragments totaling ≥500mg) was carried out by a gentleMACS dissociator system (Miltenyi) with the m_Spleen 04.01 setting, which involves churning and shredding in a sterile disposable tube. Enzymatic digestion was performed in RPMI medium at 37°C for 30 minutes. A large volume of 5% FBS/RPMI was then added to terminate the enzymatic reaction. The tissue was ground through a 70µm filter using the flat plastic end of a 3mL syringe plunger to disperse the remaining intact tissue and dispense dissociated cells.

#### Synovial biopsy tissue

For the samples obtained by synovial biopsy, tissue fragments were minced into small fragments (∼1mm^3^) using a scalpel. Samples were then subjected to enzymatic digestion at 37°C while being exposed to continuous stirring in a U-bottom polystyrene tube (12 x 75mm) with a magnetic stir bar for 30 minutes. Half way through the enzymatic digestion, samples were passed gently through a 16-gauge syringe needle ten times for additional mechanical disruption.

### RNA extraction from whole synovial tissue fragments

For samples used in the aforementioned dissociation protocol, synovial tissue fragments were also preserved for whole tissue RNA extraction. Three whole tissue replicates were collected for each sample, with six fragments placed into a cryovial containing 1mL of RNALater (Qiagen) and inverted three times. The cryovials were incubated overnight at 4°C. The next day, the cryovials were spun at ∼1000xg for 30 seconds, most of the RNAlater was removed– leaving only enough RNAlater to cover the tissue. Cryovial were then place into −70°C for storage. For RNA extraction, samples were thawed and fragments transferred into RLT lysis buffer (Qiagen) + 1% b-mercaptoethanol (Sigma) and homogenized using a TissueLyser II (Qiagen) before RNA isolation using RNeasy columns.

### Flow cytometry cell sorting

Synovial cell suspensions were stained with an 11-color flow cytometry panel designed to identify synovial stromal and leukocyte populations. Antibodies included anti-CD45-FITC (HI30), anti-CD90-PE(5E10), anti-podoplanin-PerCP/eFluor710 (NZ1.3), anti-CD3-PECy7 (UCHT1), anti-CD19-BV421 (HIB19), anti-CD14-BV510 (M5E2), anti-CD34-BV605 (4H11), anti-CD4-BV650 (RPA-T4), anti-CD8-BV711 (SK1), anti-CD31-AlexaFluor700 (WM59), anti-CD27-APC (M-T271), anti-CD235a-APC/AF750, TruStain FcX, and propidium iodide. Cells were stained in Hepes-buffered saline (20 mM HEPES, 137 mM NaCl, 3mM KCl, 1mM CaCl_2_) with 1% bovine serum albumin (BSA) for 30 minutes, then washed once, resuspended in the same buffer with propidium iodide added, vortexed briefly, and passed through a 100µm filter.

Cells were sorted on a 3-laser BD FACSAria Fusion cell sorter. Intact cells were gated according to FSC-A and SSC-A. Doublets were excluded by serial FSC-H/FSC-W and SSC-H/SSC-W gates. Non-viable cells were excluded based on propidium iodide uptake. Cells were sorted through a 100µm nozzle at 20 psi.

A serial sorting strategy was used to sequentially capture cells for bulk RNA-seq and then single cell RNA-seq if sufficient numbers of cells were present. First, 1000 cells of the targeted cell type were sorted for low-input RNA-seq into a 1.7mL Eppendorf tube containing 350uL of RLT lysis buffer (Qiagen) + 1% β-mercaptoethanol. Once 1000 cells of a particular cell type were collected, the sort was stopped and the tube was exchanged for a second tube containing FACS buffer. Sorting was then resumed and the rest of the cells of that type were collected into the second tube as viable cells. This process was carried out for four targeted populations. Live cells of each population that were sorted into FACS buffer were then re-sorted as single cells into wells of 384-well plates containing 1µL of 1% NP-40, targeting up to 144 cells of each type per sample.

### RNA sequencing

RNA from sorted bulk cell populations was isolated using RNeasy columns (Qiagen). RNA from up to 1000 cells was treated with DNase I (New England Biolabs), then concentrated using Agencourt RNAClean XP beads (Beckman Coulter). Full-length cDNA and sequencing libraries were prepared using the Smart-Seq2 protocol as previously described [16]. Libraries were sequenced on a MiSeq (Illumina) to generate 25 base pair, paired-end reads.

Single cell RNA-seq was performed using the CEL-Seq2 protocol (Illumina). Libraries were sequenced on a Hiseq 2500 (Illumina) in Rapid Run Mode to generate 76 base pair, paired-end reads. Mapping of reads to human reference transcriptome hg19 and quantification of mRNA expression levels was performed using RSEM [17]. Basic analyses of single cell RNA-seq data, including quantification of genes detected per cell, filtering of cells with less than 1000 detected genes, and principal components analyses, were performed.

### Differential expression

To examine differential gene expression profiles between RA and OA synovial tissues processed through the consensus disaggregation protocol, tissue samples were collected from 10 RA and 10 OA patients. From these 20 patients, 2-8 replicates were generated per donor for a total of 87 samples. We performed differential expression analysis by fitting a linear mixed model to each gene, in which we controlled for unwanted technical variation with donor and site as random effects.

### Gene ontology term enrichment analysis

We tested Gene Ontology (GO) enrichment with differentially expressed genes. We used Ensembl gene IDs downloaded on April 2016, including 9,797 GO terms and 15,693 genes. The minimal hypergeometric test was used to test for significance [18].

### Mass Cytometry

Synovial cells were resuspended in PBS/1%BSA with primary antibody cocktails at 1:100 dilution for 30 minutes (Additional file: Table 1). All antibodies were obtained from the Longwood Medical Area CyTOF Antibody Resource Core (Boston, MA). Cisplatin was added at 1:400 dilution for the last 5 minutes of the stain to assess viability. Cells were then washed and fixed in 1.6% paraformaldehyde for 10 minutes at room temperature. Cells were then washed and incubated with Ebioscience Transcription Factor Fix/Perm Buffer for 30 minutes, washed in Ebioscience perm buffer, and then stained for intracellular markers at 1:100 for 30 minutes. Cells were re-fixed in 1.6% paraformaldehyde for 10 minutes and stored overnight in PBS/1%BSA. The following day, cells were incubated with MaxPar Intercalator-Ir 500uM 1:4000 in PBS for 20 minutes, then washed twice with MilliQ water, filtered, and analyzed on a Helios instrument (Fluidigm). Mass cytometry data were normalized using EQ™ Four Element Calibration Beads (Fluidigm) as described [19]. viSNE analysis was performed using the Barnes-Hut SNE implementation on Cytobank (www.cytobank.org). Gated live cells (DNA-positive, cisplatin-negative) were analyzed using all available protein markers. Biaxial gating was performed using FlowJo 10.0.7.

## Results

### Synovial tissue dissociation: optimizing cell yield and surface marker preservation

Laboratories within the AMP RA network (Fig. 1a) initially set out to collectively establish a standard operating procedure for isolating cells of differing lineages from synovial tissue. This protocol required balancing rigorous dissociation of stromal cells that adhere tightly to the tissue matrix, while minimizing perturbations and maintaining viability, RNA stability and key surface proteins required for downstream isolation and assays. In addition, methods were needed to handle both large arthroplasty samples and small synovial biopsy samples (Fig. 1b).

**Figure 1.**
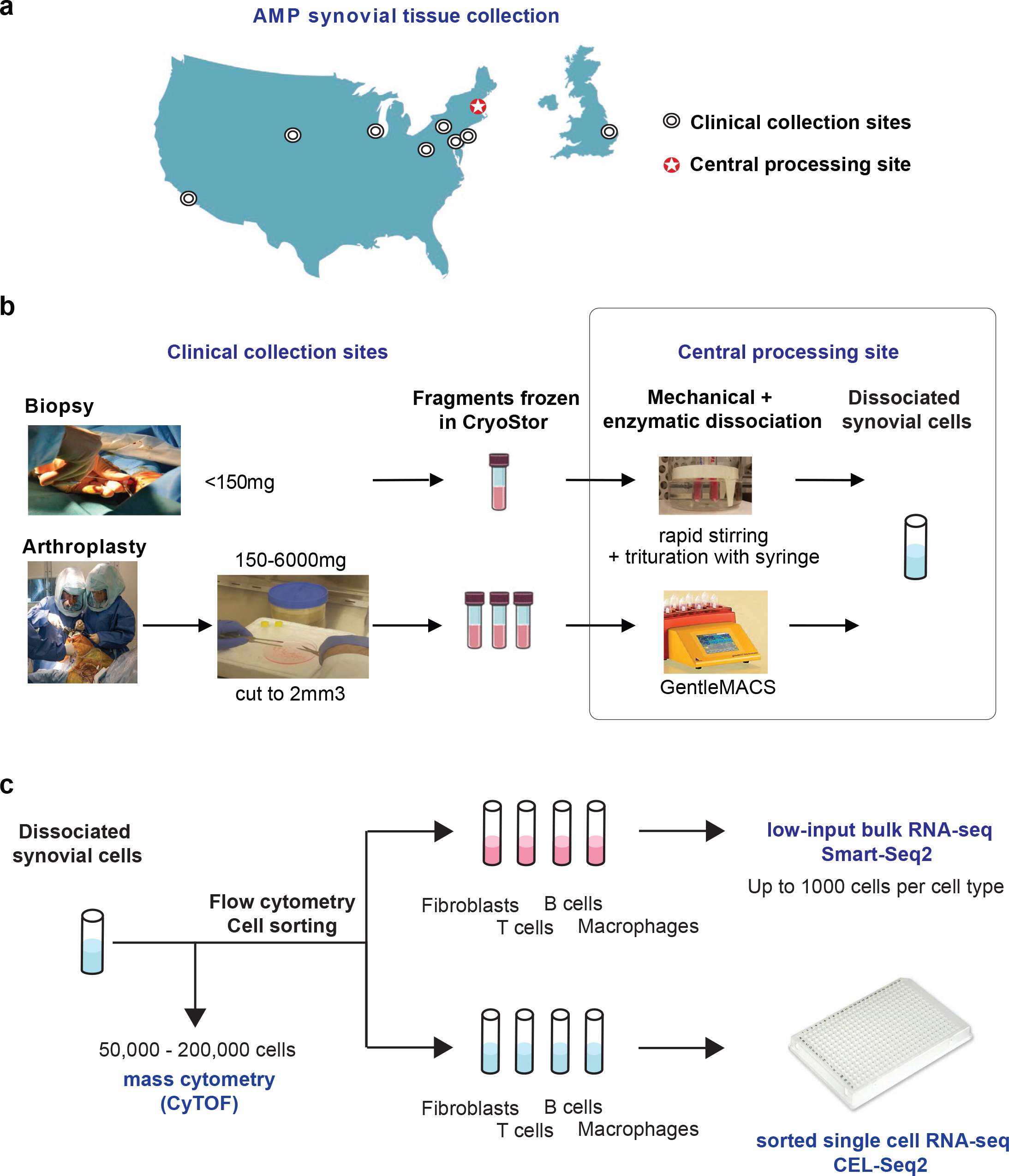
The AMP consortium pipeline for high-dimensional synovial tissue assays. a) Numerous Clinical Collection Sites contributed cryopreserved synovial samples to the Central Processing Site. b) At the Clinical Collection Sites synovial tissue was excised during research-oriented biopsies or routine removal in arthroplasty surgeries. Tissues and dissociated synovial cells were cryopreserved for centralized processing. c) Synovial tissue was mechanically disaggregated via magnetic bar stirring in a heated water bath for smaller biopsy fragments or gentleMACS dissociation for larger arthroplasty specimens. Tissue dissociation also involved incubation with protein and nucleic acid degrading enzymes at 37°C. After filtration of debris, sample aliquots were processed in parallel for mass cytometry (CyTOF) analysis and FACS-based sorting of four cell types for single-cell and bulk RNA-sequencing.

### Selection of methods for mechanical disruption

Pilot experiments across several sites suggested that various forms of mechanical disruption could be used, including manual inversion of the tube or incubation in a Stomacher^®^ 400 circulator. Dissociation using the GentleMACS was selected for arthroplasty as this method provided an automated and standardized protocol that could be implemented across laboratories (Fig. 1c). However, dissociation of small synovial biopsy fragments (∼0.5-1mm^3^) using the GentleMACS system generated variable and insufficient yields, possibly due to cell retention in the dissociation container. Therefore, synovial biopsy samples were subjected to mechanical disruption by continuous magnetic stirring, combined with trituration through a syringe needle.

### Enzymatic dissociation combined with mechanical disruption generates high cell yields

To test the impact of combining enzymatic treatment with mechanical dissociation, proteolytic enzymes were added to the media during mechanical dissociation. For the majority of tissue samples, higher cell yields were obtained when synovial fragments were incubated with enzyme in addition to mechanical disruption (Fig. 2a). Importantly for highly adherent stromal (PDPN+CD45-) lineages, the proportion recovered increased with enzymatic digestion (Fig. 2b, c).

**Figure 2.**
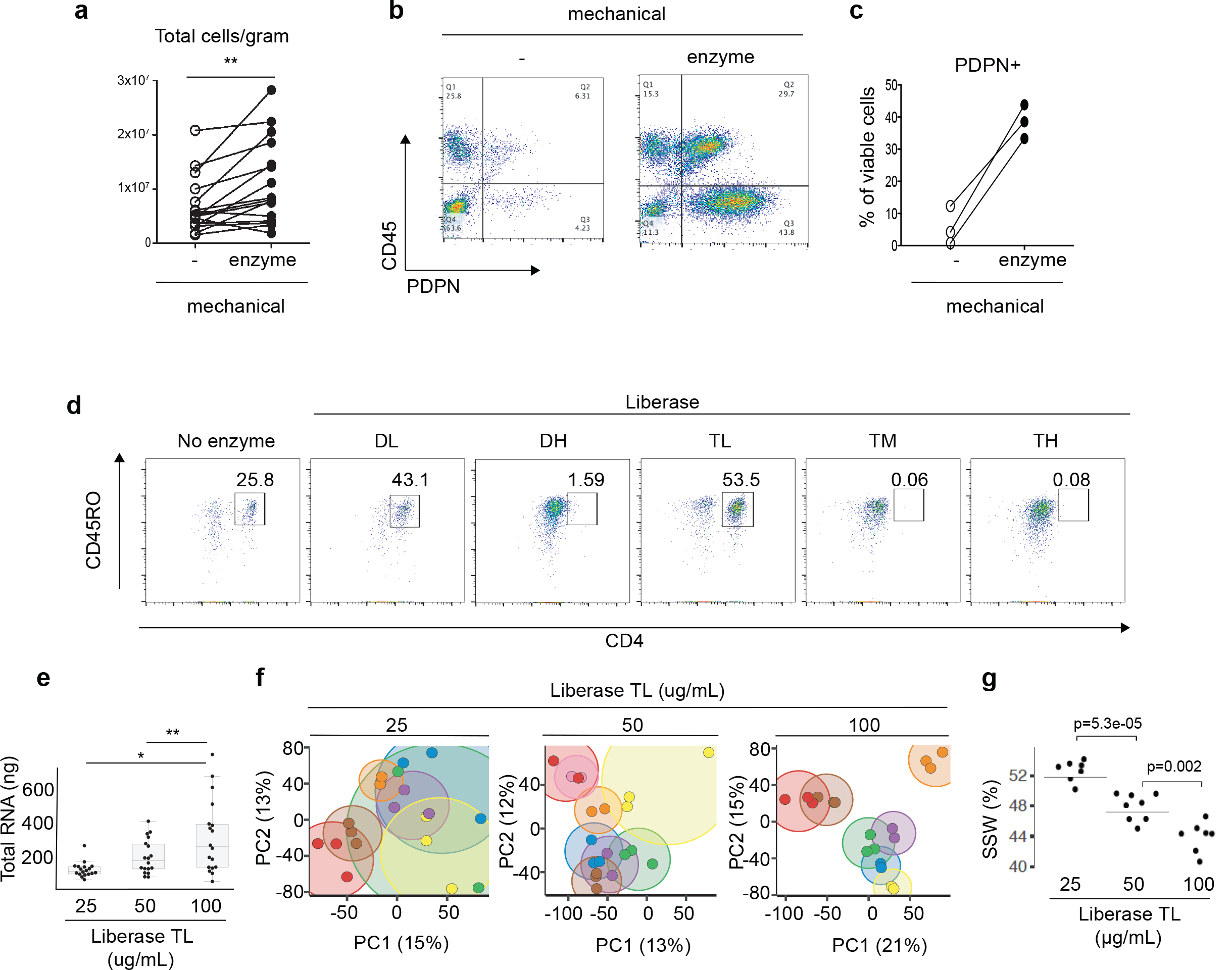
High synovial cell yield, preserved surface markers and reproducible transcriptomic results with a mechanical and enzymatic disaggregation protocol. a) Total cell counts per gram from synovial tissue mechanically disaggregated with or without Liberase TL proteolytic enzyme treatment. n=16, paired T test. b-c) Flow cytometry detection and quantification of stromal cells (CD45-PDPN+) upon mechanical disaggregation with or without Liberase TL proteolytic enzyme treatment. PDPN, podoplanin. d) Flow cytometry of dissociated synovial cells treated with a panel of proteolytic Liberase formulations. Cells were gated for viability and CD3 T cell receptor subunit. Plots are representative of n=4 biologic replicates. e) Total RNA yield from synovial tissue dissociated with three concentrations of Liberase TL. Two or three technical replicates from nine tissues were used for each enzyme concentration. ANOVA. f) Principal component analyses on the RNA-seq gene expression results for the samples from e, whereby each technical replicate of the same sample is represented by a dot with the same color while the variation between the replicates is indicated by the size of the encircling cloud of the same color. g) Sum of squares within (SSW) replicates from the same donor divided by the total sum of squares, using the gene expression profiles from e and f. Horizontal lines indicate the SSW values using all replicates. The individual dots are SSW values computed after a leave-one-out strategy where replicates from one donor are left out and SSW is computed with the remaining samples. Student’s t test was used to test if the bootstrapped SSW values differ significantly.

### Proteolytic enzyme preparations differ in surface marker preservation

Focusing on highly purified enzyme formulations designed for standardized protocols, the consortium tested Liberase^™^ mixtures containing thermolysin (T) or dispase (D) in combination with collagenase. Across the enzyme panel, we focused on identifying an optimized enzymatic formulation that preserved important cell surface proteins such as CD3 and CD4. Certain formulations compromised the detection of cell surface proteins. For example, on CD3+ T cells isolated from synovium, a decline in CD4 surface detection was observed with Liberase Dispase-High (DH), Thermolysin-Medium (TM) and Thermolysin-High (TH) compared to no enzyme treatment or Thermolysin-Low (TL) and Dispase-Low (DL) (Fig. 2d). As peripheral blood mononuclear cell preparations with equal numbers of circulating T cells resulted in a similar pattern of loss in CD4 detection when treated with the enzymes, the decline in CD4 levels in the synovial preparations is likely due to direct degradation of the cell surface protein by proteases rather than a selective reduction in extracting CD4+ T subsets (Additional file: Supplementary Fig. 1a). As Thermolysin-low (TL) versus TM Liberase mixtures proved equally efficient at dissociating out both stromal and hematopoietic cells (Additional file: Supplementary Fig. 1b), Liberase TL^™^ was chosen for both high cell yields and preservation of cell surface markers for downstream analyses, including mass cytometry.

### Optimization of proteolytic enzyme concentrations

To determine the optimal concentration of proteolytic enzyme for consistent cell recovery and reproducibility of downstream assays, four Network sites processed arthroplasty samples using three different Liberase TL^™^ concentrations, with three technical replicates for each concentration. In comparison to the 25 and 50µg/ml concentrations, the 100µg/ml concentration yielded the highest amount of intact RNA after tissue dissociation (Fig 2e). In addition, synovial samples processed with 100µg/mL enzyme concentration demonstrated the lowest variance in

RNA-seq gene expression across replicate samples, visually depicted in the principal component analysis (PCA) plots where distance between replicates is minimized with the 100µg/mL concentration (Fig. 2f). Next, we computed the sum of squares within (SSW) to quantify replicate similarities within each of the three enzyme concentrations. Here, we also observed replicates processed with 100µg/mL had the least variation (Fig. 2g).

To gain insight into the source of gene expression variation across synovial samples, we applied an analysis of variance (ANOVA) to each principal component to assess the contribution of the following sources: site, donor, and the potential impact of the dissociation process (Additional file: Supplementary Fig.1c). The majority of gene expression variation associated with donor differences (PC1 and 2), which encompasses biological differences such as disease state. Site-specific collection and/or processing differences also contributed variation (PC1). Of note, the AMP network has designated a central processing site to eliminate site-specific dissociation effects.

Sample variability in PC3 and PC6 also related to gene expression differences that were identified in a separate RNAseq analysis comparing whole versus dissociated synovial preparations (Additional file: Supplementary Fig. 1c-e). The dissociated versus whole tissue analysis was intended to identify dissociation-induced effects, however, involved mixtures of various cell types and, correspondingly, much of the gene expression differences indicated cell composition differences. These differences likely relate to differences in the preparation protocols; for example, adipocytes escape during spinning and aspiration steps in the dissociation protocol, while red blood cells may inadvertently be removed in whole tissue preparations during a salt solution incubation and subsequent aspiration. Nonetheless, outside of cell type-specific genes, there was an increase in stress response genes in dissociated cell preparations, similar to a previous report on dissociation-induced effects in single-cells [20].

This included an upregulation in heat shock protein genes (*HSP1A1*, *HSP6A*, *HSP90AA*) and early response genes (*FOS*, *JUN*, *ATF3*) (Supplemental Fig. 1d-e). Thus, synovial cells demonstrating specific upregulation in this signature could indicate a dissociation-induced gene expression pattern rather than a cell subset-specific program. For subsequent single-cell RNAseq studies, individual synovial cells with high levels of this stress response could either be removed from the analyses or the signature could be adjusted for computationally.

Overall, these results suggest that dissociation with 100µg/mL Liberase TL produces consistent and robust recovery of diverse synovial stromal and immune cell populations with the largest source of variation in the samples due to donor biological differences.

### Development of Cryopreservation methods

Having established synovial dissociation methods, we then focused on establishing a strategy to analyze samples acquired at distant sites through a uniform high-dimensional analysis pipeline. We first evaluated the feasibility of studying synovial cells cryopreserved after tissue dissociation. From three different arthroplasty samples analyzed either immediately after dissociation (fresh) or after freezing the cells in a cryopreservation solution for at least one day, comparable cell frequencies were extracted for synovial fibroblasts, monocytes, T cells, and B cells (Figure 3a). This result suggested that synovial cells generally survive cryopreservation well, consistent with prior studies [5, 21, 22].

**Figure 3.**
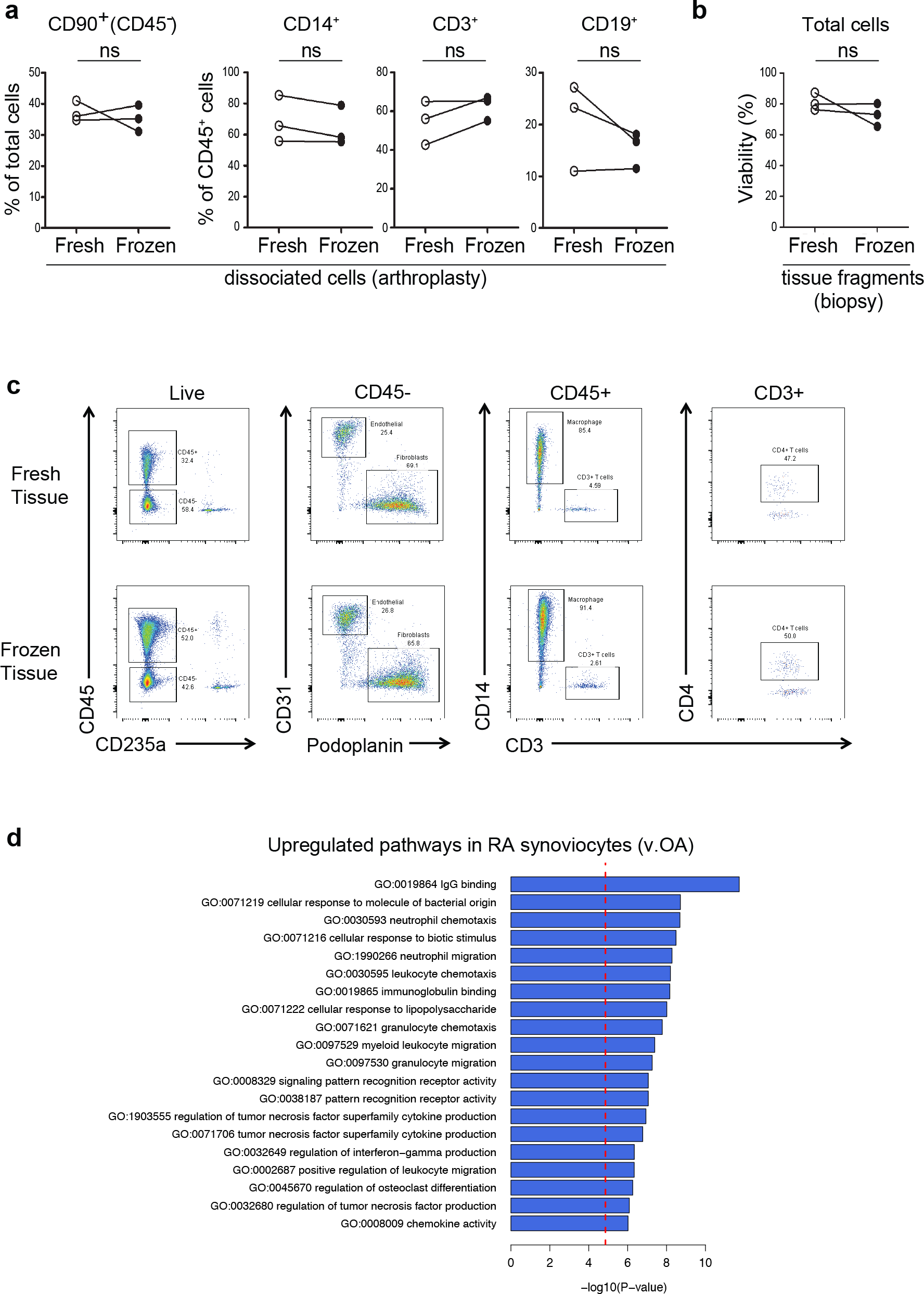
Cryopreservation of synovial fragments retains stromal and hematopoietic cell viability and RA synovial gene expression patterns. a) Frequency of synovial cell subpopulations from arthroplasty samples dissociated from freshly isolated or cryopreserved aliquots. ns, no significant difference. b) Cell viability frequency assayed by flow cytometry for synovial biopsy samples dissociated from fresh or cryopreserved tissue fragments. c) Flow cytometry analysis of cells dissociated from synovial tissue either immediately or after cryopreservation. d) Bar plot showing the top 20 GO terms enriched for genes upregulated in RA patient disaggregated synovial cells in comparison to patients with osteoarthritis (OA).

In addition to cryopreserving dissociated synovial cells, we next evaluated the feasibility of cryopreserving intact synovial tissue fragments prior to dissociation. Notably, synovial biopsy tissue processed after cryopreservation yielded cells with comparable viability to paired tissue analyzed immediately after isolation (Fig. 3b). In addition, flow cytometric analysis of cells from cryopreserved synovial tissue showed robust detection of stromal and hematopoietic cell populations, similar to fresh synovial tissue (Fig. 3c). These results indicate that intact synovial tissue can be cryopreserved for subsequent cellular analyses.

We next examined whether application of the protocol implementing 100µg/mL of Liberase TL, mechanical disaggregation and viable freezing could detect robust expression pattern differences between dissociated synovial cells from tissues from patients with RA and osteoarthritis (OA). Importantly, an RNA-seq analysis of disaggregated total synovial cells demonstrated an up-regulation in RA of genes involved in immune processes, such as immunoglobulin binding, cellular response to molecules of bacterial origin, leukocyte chemotaxis and migration, cytokine production and antigen processing and presentation by MHC class II (Fig. 3d).

### High-dimensional assays of synovial cells

A goal of the AMP Network is to generate both mass cytometry and RNA-seq analyses of cells from cryopreserved synovial samples dissociated by the protocol described above. Acquisition of data for both of these technologies from the same synovial sample allows the coupling of detailed quantification of cell populations with genome-wide transcriptomic analyses. These combined analyses enable interrogation of correlations between cell populations and transcriptomic signatures with unprecedented resolution for a human autoimmune disease tissue target. Because synovial samples vary widely in both cell yield and composition, we developed an algorithm to allocate cells for different analyses in a step-wise fashion without prior knowledge of the cell composition (Figure 1c). Preliminary experiments indicated that reproducible mass cytometry analyses could be obtained from 100,000 synovial cells; therefore, for samples that yielded over 200,000 synovial cells, approximately half of the cells were allocated to mass cytometry, with the rest were used for sequential low-input and single cell RNA-seq analyses. When less than 200,000 synovial cells were obtained, mass cytometric analysis was omitted and only RNA-seq analyses were performed.

### Mass cytometry of synovial cells

A 35-parameter mass cytometry panel was developed to identify stromal, vascular, and immune cells within synovial samples (Figure 4a). Visualization of the mass cytometry data obtained from a cryopreserved synovial biopsy sample using the dimensional reduction tool viSNE [23] revealed clear discrimination of synovial fibroblasts (cadherin-11+), endothelial cells (VE-cadherin+), and immune cell subsets from cryopreserved RA synovial tissue (Figure 4b). viSNE visualization highlighted the heterogeneity within immune cell populations, which was confirmed by biaxial gating. For example, two populations of B cells were easily resolved, including one CD19+ CD20+ population, and a distinct CD19+ CD20-CD38^hi^ population comprised of antibody-secreting cells (Figure 4b, c) [24, 25]. Marked heterogeneity within the CD3+ T cell population was also evident, with multiple lobes of T cells visualized, including a subset of PD-1^hi^ ICOS+ CD4+ T cells (Figure 4b, c) [5]. This visualization also demonstrated heterogeneity among the podoplanin+ synovial fibroblasts, including fibroblast subpopulations that express CD90 and/or CD34 [21, 22]. These results indicate that high-resolution mass cytometry data can be obtained from synovial cells collected from cryopreserved intact synovial tissue that is subsequently thawed and dissociated.

**Figure 4.**
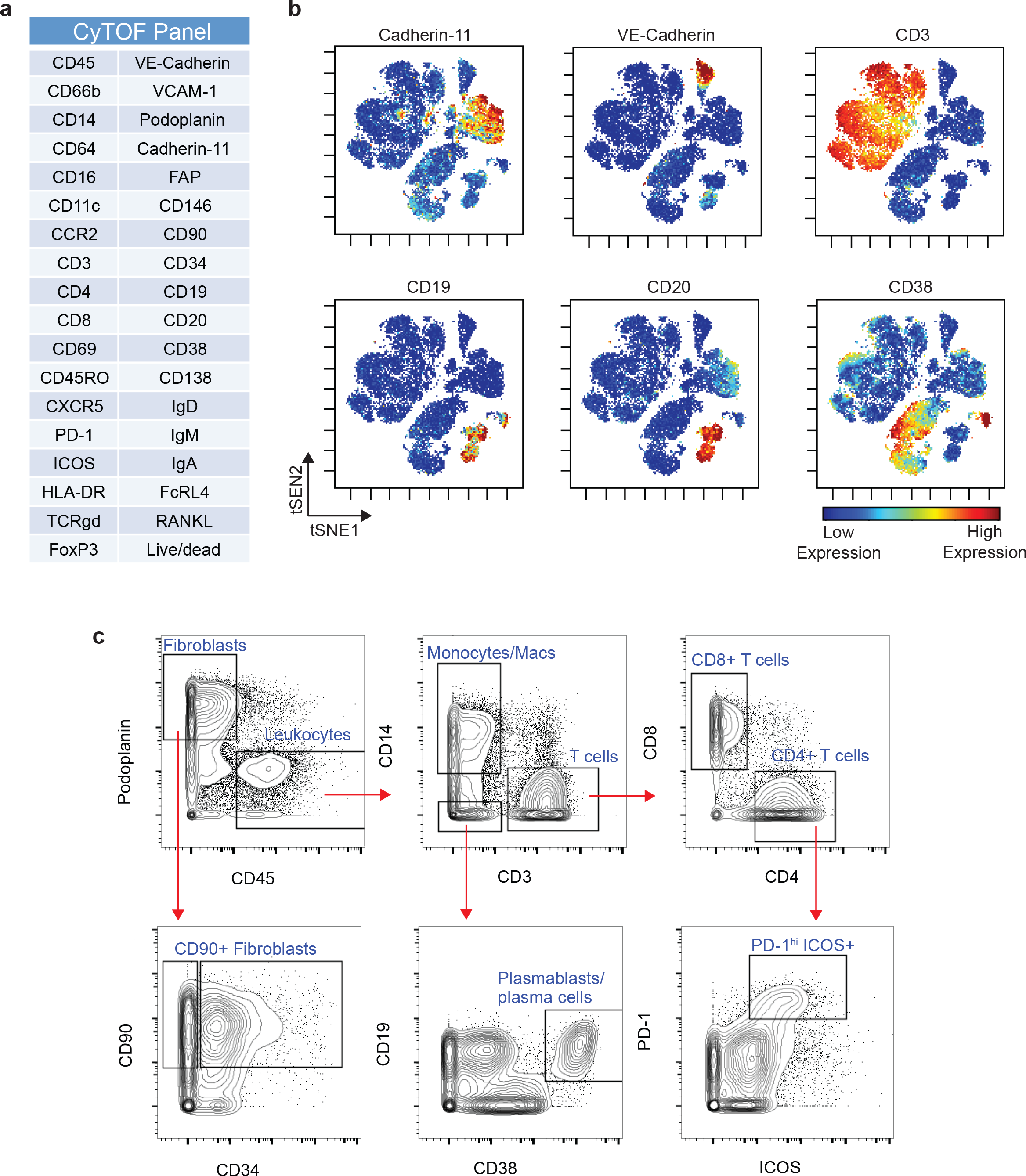
Mass cytometry analysis of dissociated synovial tissue reveals synovial cell heterogeneity via surface marker detection. a) The antibody marker panel for synovial cell mass cytometry analysis. b) viSNE visualization of synovial cells analyzed by mass cytometry from a cryopreserved RA synovial biopsy sample. Each dot is a cell. The color reflects the level of expression of the marker indicated at the top of the plot. c) Biaxial contour plots of the same mass cytometry data shown in (b), showing serial gating of cell subpopulations.

### RNA-seq transcriptomics of synovial cell populations and single-cells

In addition to mass cytometry analyses, a major goal of the AMP Network is to define molecular signatures and pathways of disease by transcriptome analysis. Low-input RNA-seq provided robust gene expression values for a large number of protein coding genes averaged across approximately 1000 cells. The complementary single-cell RNA-seq approach provided a high-resolution view of the heterogeneity of expression profiles between individual cells. We designed a cell sorting workflow that prioritized collecting cells for low-input RNA-seq, with a goal of collecting 1000 cells from each of the four targeted cell types. If more than 1000 cells of a targeted cell type were available, the remaining cells were sorted into a second collection tube for subsequent single-cell RNA-seq analysis (Figure 1c). This sorting scheme enabled capture of virtually all of the cells of each of the four target populations for low-input and single cell RNA-seq.

Synovial cells were stained with a multidimensional flow cytometry panel that could identify diverse cell populations including fibroblasts (CD45^-^ podoplanin^+^ CD31^-^), endothelial cells (CD45^-^ podoplanin^-^ CD31^+^), macrophages (CD45^+^ CD14^+^ CD3^-^) and T cells (CD45^+^ CD14^-^ CD3^+^) (Figure 5a). Low-input RNA-seq analysis of fibroblasts, macrophages, endothelial cells, and T cells sorted from three cryopreserved RA synovial biopsy samples yielded transcriptomes with 8,000-13,000 genes detected from cell inputs ranging from 1000 cells down to 83 cells (Figure 5a, b). Principal components analysis robustly distinguished the different cell types at the global transcriptomic level (Figure 5c). These results indicate that cells obtained from cryopreserved biopsies yield transcriptomes that reflect the expected cell identity.

**Figure 5.**
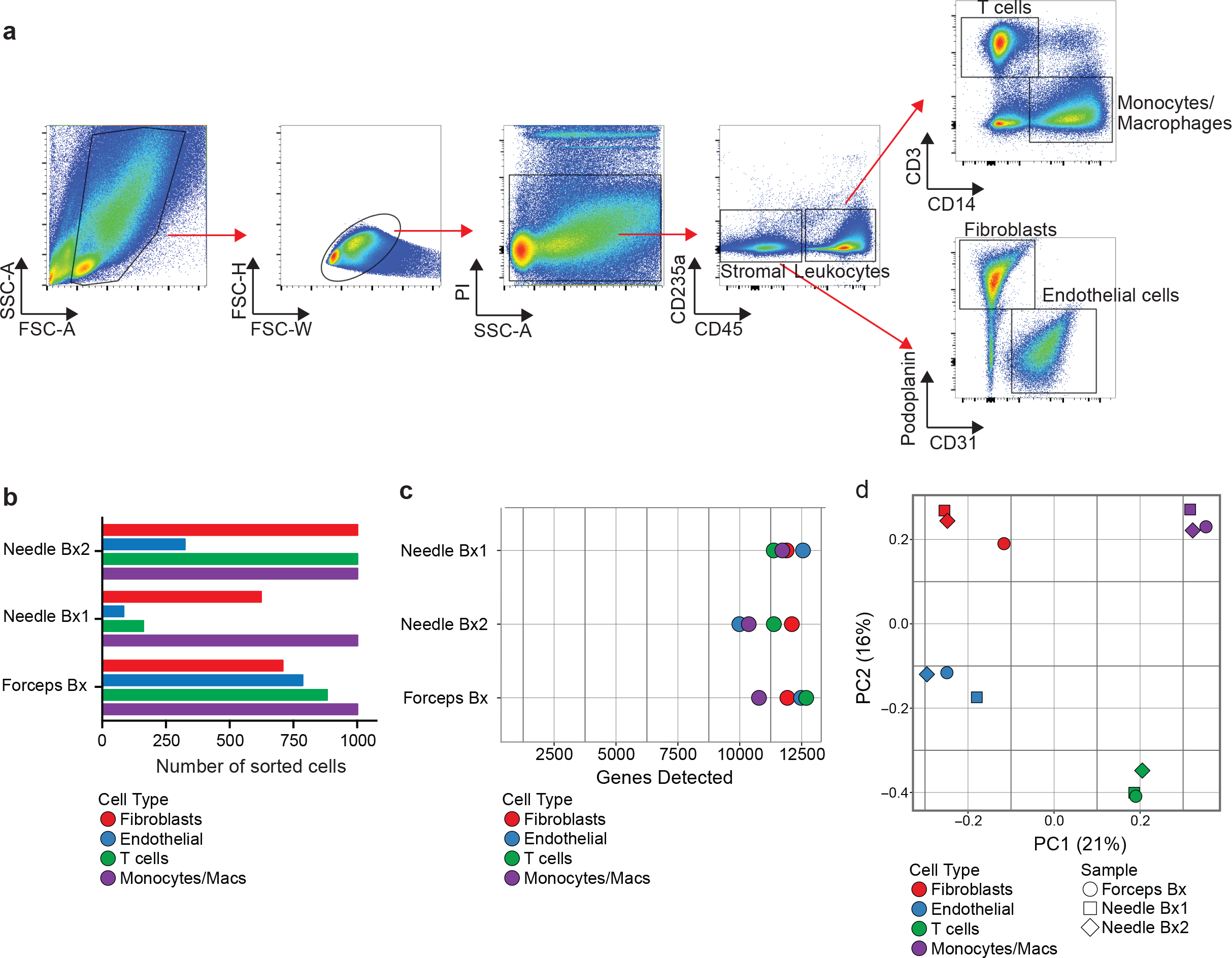
Low-input RNA-seq distinguishes cell populations sorted from dissociated RA synovial cells. a) Flow cytometric detection of distinct cell populations obtained from a cryopreserved RA synovial biopsy. b) Number of cells of each targeted cell populations sorted from 3 different RA synovial biopsy (Bx) samples (2 needle biopsies, 1 portal and forceps) for low-input RNA-seq. c) Number of genes detected by low-input RNA-seq for each cell population sorted from the same biopsies as in (b). d) Principal components analysis of sorted cell populations from the 3 RA synovial biopsy samples based on the 5006 genes with greatest variance. Color indicates the sorted cell type, while shape indicates the individual donor.

Transcriptomes of single cells sorted from cryopreserved RA synovium were obtained by RNA-seq through the CEL-Seq2 platform. High quality transcriptomes from both from single cell fibroblasts and leukocytes from RA synovium were generated, with fibroblasts generally yielding higher gene counts (Figure 6a). Principal components analysis of single cells from an RA synovial sample separated the fibroblasts from the immune cells along PC1, and mostly separated macrophages from T and B lymphocytes along PC2 (Fig. 6b). Importantly, single cells from each sorted cell population expressed markers expected for that cell type. For example, the *podoplanin* transcript was expressed primarily in fibroblasts, consistent with flow cytometry and mass cytometry data, while the *CD14* transcript was identified in sorted macrophages, an expected finding since the cells were flow sorted by CD14 protein expression (Fig.6c). Similarly, the *CD3E* transcript was uniquely detected in sorted T cells, while *CD79A* expression was unique to sorted B cells. These results indicate that high-quality transcriptomes can be generated from bulk populations and single cells obtained from cryopreserved RA synovial tissue samples.

**Figure 6.**
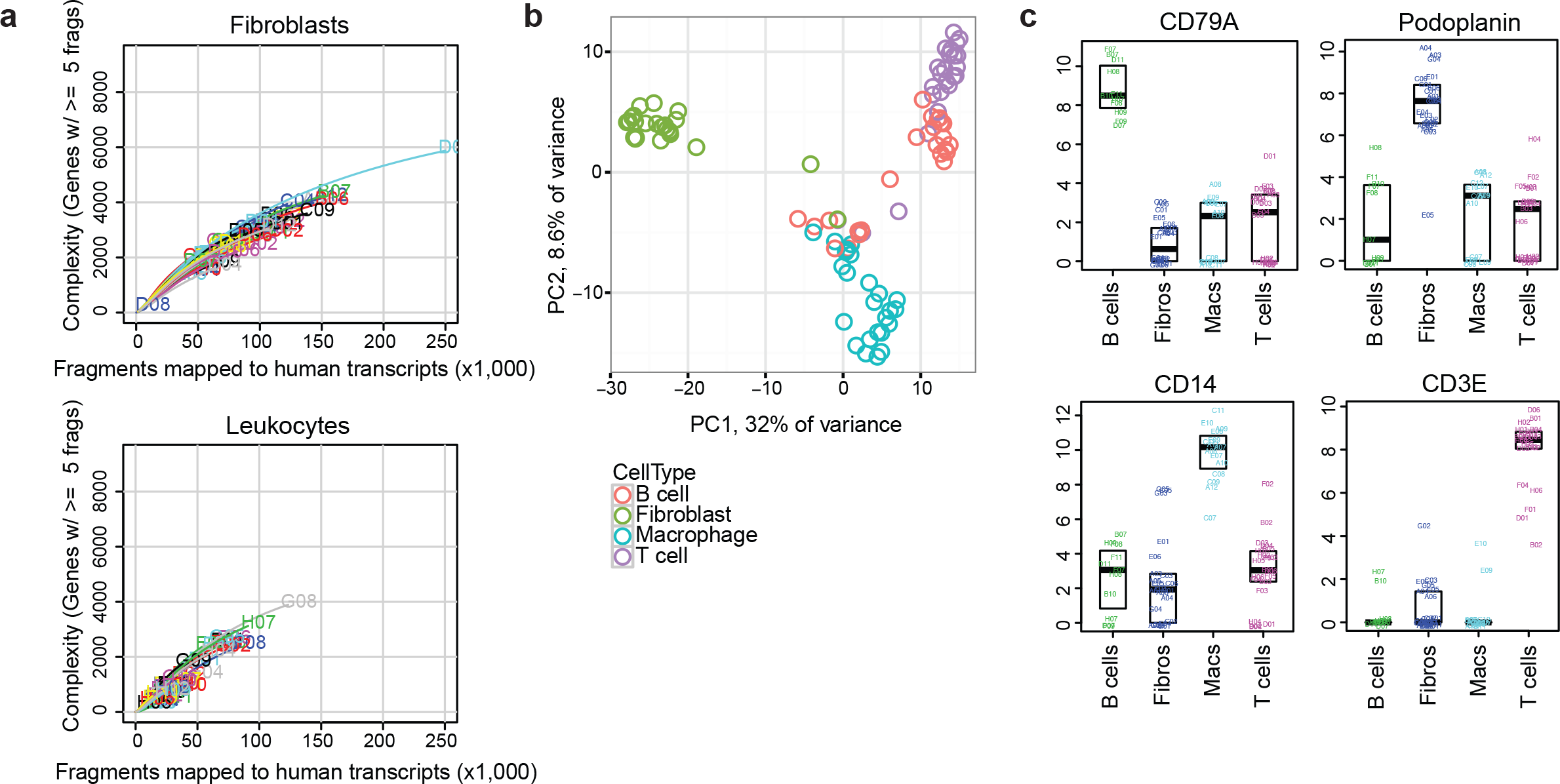
Single-cell RNA-seq transcriptomic analysis distinguishes different cell types sorted from cryopreserved RA synovium. a) Saturation curves indicating the complexity of transcriptomes obtained from single cells (fibroblasts or leukocytes) sorted from cryopreserved RA arthroplasty tissue. At least five read fragments were required for a gene to be considered “detected”. b) Principal components analysis of single cells from cryopreserved RA synovium, colored by cell type as determined by flow cytometry at the time of cell sorting. The top 648 most variable expressed genes (F statistic) were used, centered and scaled. Plotted are PC scores for PC1 (x-axis) and PC2 (y-axis). c) Expression (log2(TPM+1)) of selected lineage-characteristic genes in sorted cell populations from a cryopreserved RA arthroplasty sample. A total of 96 cells (24 for each type) were filtered down to 69 by eliminating low-quality cells (those with more than 10% of reads in the well mapped to ERCC) and presumed doublets (those with greater than 2.8 million reads).

## Discussion

Here we describe the development of an experimental pipeline to analyze synovial tissue with multiple high-dimensional technologies, aimed at deconstructing the cellular and molecular features of RA. This standardized and validated strategy enables single-cell analytics on large sample numbers collected from numerous clinical sites.

The integration of contemporary high-dimensional assays provides the opportunity to dissect the properties of tissues affected by rheumatic disease with a resolution not previously possible. Histologic and immunohistochemical assessments of synovium have revealed marked heterogeneity in RA synovial pathology [13, 26]. Gene expression analyses of synovium by microarray have added gene expression signatures to these histologic patterns [27–30]. However, broad analyses of gene expression in synovium have thus far been limited mainly to assessments of whole tissue, which cannot distinguish which cell types express the relevant genes [27, 31–34]. The analysis pipeline described here enables simultaneous quantification of a wide range of stromal and leukocyte subpopulations by mass cytometry, paired with genome-wide transcriptional analyses, both of targeted cell populations and single cells. Mass cytometry alone provides a powerful, broad assessment of the cell types and phenotypes within the tissue on a relatively large number of cells [5]. Transcriptomic analyses of sorted subpopulations can identify the cell types contributing specific gene expression signatures derived in the synovium, including signatures specific to RA synovium. Single cell transcriptomics allow a high-powered characterization of the heterogeneity of cells across a tissue [35] and within cell subsets that were previously classified as the same cell type [21, 22, 36]. We propose that integrating the multiple high-dimensional datasets acquired with this pipeline will reveal how distinct cell types contribute to specific aspects of RA pathology. Applied to a large patient cohort, these analyses have the potential to reveal functional pathologic markers that distinguish patients with distinct clinical trajectories and therapeutic responses [5].

Implementing these technologies requires robust tissue processing protocols. The validation studies described here, which were carried out by a collaboration among laboratories with expertise in synovial tissue and RA, provide a widely applicable, reproducible method for obtaining single cells from RA synovium that performs well in cytometric and transcriptomic analyses. Classically, synovial tissue has been disaggregated by crude collagenase preparations, which often cleave cell surface proteins and may contain bacterial products that stimulate immune cells. Here, varied concentrations of highly-purified proteolytic enzyme preparations were assessed for preservation of vulnerable cell surface proteins required for downstream assays, as well as cell viability, RNA integrity, and applicability to next-generation sequencing. Combining limited enzyme incubation time with mechanical disaggregation yielded viable cell populations from disparate lineages that performed well in downstream assays. It is important to recognize that dissociating tissues into cell suspensions may introduce processing signatures [20]. However, differences in transcriptome between intact and dissociated tissue may arise from a number of sources. These may include better representation of genes from cells difficult to extract in the intact tissue samples, in addition to potentially adverse consequences such as a cellular response to tissue manipulation. Importantly, with the synovial tissue disaggregation protocols developed here mass cytometry of the single cells obtained clearly identified multiple expected cell populations in striking detail. RNA-seq transcriptomic analyses also robustly distinguished cells of different types at both the low-input and single cell level. Further, transcriptomic analysis of synovial cells identified RA-specific gene signatures even in the relatively small sample set presented with the largest source of variation in the transcriptome due to donor biological differences. While dissociation-induced gene signatures are identifiable, they can be adjusted for computationally if necessary. Taken together, these observations strongly argue for the potential of this approach to discover new aspects of RA cellular pathology, which can be further validated in intact tissues.

The ability to effectively cryopreserve synovial tissue marks a major technical advance for synovial research. The viability and flow cytometric phenotypes of cells from cryopreserved synovial tissue samples was comparable to those from freshly processed tissues. Cells obtained from cryopreserved tissue samples were largely viable and readily used for all of the mass cytometric and RNA-seq analyses presented in this report. Cryopreservation of intact synovial tissue provides a simple, rapid technique to preserve and transport affected tissues for cellular studies in a multi-site consortium. Tissue fragment cryopreservation enables sample collection from numerous clinical sites, including those lacking the resources for tissue dissociation. This approach also eliminates site-specific processing effects, allowing samples to be processed at a central site that possesses multiple high-dimensional technologies, and to utilize batch processing of samples to minimize technical variation and confounding effects.

In an ongoing Phase 1 study, the AMP RA Network has implemented this synovial tissue pipeline in a pilot cohort of over 50 RA and OA patients with the goal of defining RA disease-specific cellular populations and pathways. This will be followed by a Phase II study involving a much larger cohort of RA patients, including those with early disease, with a focus on identifying RA tissue biomarkers and novel tissue drug targets. A similar strategy of tissue cryopreservation has been adopted by the AMP for single cell transcriptomic analyses of kidney biopsy samples from patients with lupus nephritis, including development of a dissociation protocol optimized for human kidney tissue [37]. Although different tissues and cellular subsets may respond to cryopreservation differently, the strategy of analyzing cells from cryopreserved tissue appears readily adaptable to multi-center studies of tissues in other inflammatory diseases.

## Conclusions

These studies demonstrate the feasibility and potential of analyzing viable cells from cryopreserved synovial tissue samples by multiple, complementary high-dimensional analyses. Using an optimized dissociation protocol that provides high yields of viable cells with preserved cell surface and transcriptomic features, synovial tissue samples acquired across a multi-site network can be analyzed in a uniform way in order to identify dominant cell types and pathways, as well as to characterize previously unidentified cellular heterogeneity. Such approaches applied to large numbers of patients offer new opportunities to discover rheumatic disease biomarkers, targets for drug development, and molecular stratification of synovial pathology.

### Ethics approval and consent to participate

The study received institutional review board approval at each site. Sites performing research synovial biopsies included informed consent specifically for this procedure. Where samples were used for transcriptome analysis, informed consent included consent for genetic analysis and deposition of AMP project data into public NIH databases.

### Consent for publication

Not applicable

### Availability of data and material

In addition to the data included in the manuscript, additional datasets analyzed during the current study are available from the corresponding author on reasonable request. Select datasets are available at the ImmPort repository.

### Funding

This work was supported by the Accelerating Medicines Partnership (AMP) in Rheumatoid Arthritis and Lupus Network (AMP RA/SLE Network). AMP is a public-private partnership (AbbVie Inc., Arthritis Foundation, Bristol-Myers Squibb Company, Lupus Foundation of America, Lupus Research Alliance, Merck Sharp & Dohme Corp., National Institutes of Health, Pfizer Inc., Rheumatology Research Foundation, Sanofi and Takeda Pharmaceuticals International, Inc.) created to develop new ways of identifying and validating promising biological targets for diagnostics and drug development Funding was provided through grants from the National Institutes of Health (UH2-AR067676, UH2-AR067677, UH2-AR067679, UH2-AR067681, UH2-AR067685, UH2-AR067688, UH2-AR067689, UH2-AR067690, UH2-AR067691, UH2-AR067694, and UM2-AR067678). This report includes independent research supported by the National Institute for Health Research/Wellcome Trust Clinical Research Facility at University Hospitals Birmingham NHS Foundation Trust. The views expressed in this publication are those of the author(s) and not necessarily those of the NHS, the National Institute for Health Research or the Department of Health. Funding was also provided by Arthritis Research UK (Fellowship 18547 and the RACE Rheumatoid Arthritis Pathogenesis Centre of Excellence grant 20298).

### Authors’ contributions

**Tissue collection, acquisition of data, data interpretation:** LTD, MJM, JDT, NM, KM, JH, SK, SMG, CP, EMG, LWM, VPB, AF, DLB, JHA. **Tissue processing and validation studies, acquisition of data, data interpretation:** LTD, DAR, KW, MJM, JDT, NM, FM, JK, KM, JH, CR, ER, SK, ABP, JAL, LBI, CP, GSF, LWM, AF, DLB, MBB, JHA. **AMP Tissue Working Group and Experimental Design:** LTD, MJM, JDT, NM, FM, KM, JH, AF, DLB, MBB, JHA. **RNA purification, library prep and sequencing:** DAR, KW, FM, DJL, JK, TME, SL, EPB, CN, NH. **Computational data analysis:** KS, MG,,DJL, TME, SL, EPB, CN, JAL, NH, SR, MBB. **Mass Cytometry experimental design, data acquisition, data interpretation:** DAR, JAL. **AMP Systems Biology Group:** KS, MG, DJL, TME, SL, EPB, WHR, NH, SR, MBB**. Manuscript Drafting:** LTD, DAR, AF, MBB, JHA**. Manuscript Review:** all. **AMP RA Disease Focus Group and Experimental Design:** EMG, PKG, LWM, VPB, MBB, JHA. **AMP Principal Investigators:** MJM, ABP, LBI, WHR, PJU, EMG, BFB, JAL, NH, CP, PKG, GSF, SR, LWM, VMH, VPB, AF, DLB, MBB, JHA.

### Accelerating Medicine Partnership Consortium; additional members

Jane Buckner, Derrick Todd, Michael Weisman, Ami Ben Artzi, Lindsy Forbess, Joan Bathon, John Carrino, Oganna Nwawka, Eric Matteson, Robert Darnell, Dana Orange, Rahul Satija, Diane Horowitz, Harris Perlman, Art Mandelin, Louis Bridges, Laura B. Hughes, Arnold Ceponis, Peter Lowry, Paul Emery, Ahmed Zayat, Amir Aslaam, Karen Salomon-Escoto, David Fox, Robert Ike, Andy Cordle, Aaron Wise, John Ashton, Javier Rangel-Moreno, Christopher Ritchlin, Darren Tabechian, Ralf Thiele, Deborah Parks, John Akinson, Chiam Putterman, Evan Der, Elena Massarotti, Michael Weisman, David Hildeman, Richard Furie, Betty Diamond, Michelle Petri, Diane Kamen, Melissa Cunningham, Jill Buyon, Iris Lee, Hasan Salameh, Maureen McMahon, Ken Kalunian, Maria Dall’Era, David Wofsy, Mattias Kretzler, Celine Berthier, William McCune, Ruba Kado, Wiliam Pendergraft, Dia Waguespack, William Apruzzese, Yanyan Liu, Gerald Watts, Arnon Arazi, Rohit Gupta, Holden Maecker, Patrick Dunn, Rong Mao, Mina Pichavant, Quan Chen, John Peyman, Ellen Goldmuntz, Justine Buschman, Jennifer Chi, Su-Yau Mao, Susana Serrate-Sztein, Yan Wang, Thomas Tuschl, Yvonne Lee, Chamith Fonseka, Fan Zhang, Ilya Korsunskiy, Judith A. James, Joel Guthridge.

## Acknowledgements

We acknowledge the many clinical coordinators and patients who helped to provide samples for analysis in this study. We appreciate the many efforts of Mina Pichavant Clinical Research Project Manager for AMP and Bill Apruzzese Associate Director of Operations and Management for AMP.

