## Supplementary Materials for "High dimensional analyses of cells dissociated from cryopreserved synovial tissue"

| Metal | Target | Clone |
|---|---|---|
| 141Pr | CD45 | HI30 |
| 142Nd | CD19 | HIB19 |
| 143Nd | RANKL | MIH24 |
| 144Nd | CD64 | 10.1 |
| 145Nd | CD16 | 3G8 |
| 146Nd | CD8α | RPA T8 |
| 147Sm | FAP | Poly |
| 148Nd | CD20 | 2H7 |
| 149Sm | CD45RO | UCHL1 |
| 150Nd | CD38 | HIT2 |
| 151Eu | PD-1 | EH12.2H7 |
| 152Sm | CD14 | M5E2 |
| 153Eu | CD69 | FN50 |
| 154Sm | CXCR5 | J252D4 |
| 155Gd | CD4 | RPA T4 |
| 156Gd | Podoplanin | NC-08 |
| 158Gd | CD3 | UCHT1 |
| 159Tb | CD11c | Bu15 |
| 160Gd | FcRL4 | 413D12 |
| 161Dy | CD138 | MI15 |
| 162Dy | CD90 | 5E 10 |
| 163Dy | CCR2 | K036C2 |
| 164Dy | Cadherin11 | 23C6 |
| 165Ho | FoxP3 | PCH101 |
| 166Er | CD34 | 581 |
| 167Er | CD146 | SHM-57 |
| 168Er | IgA | 9H9H11 |
| 169Tm | TCRgd | B1 |
| 170Er | ICOS | C398.4A |
| 171Yb | CD66b | G10F5 |
| 172Yb | IgM | MHM-88 |
| 173Yb | CD144 | BV9 |
| 174Yb | HLA-DR | L243 |
| 175Lu | IgD | IA6-2 |
| 176Yb | VCAM-1 | STA |
| 195Pt | Live/Dead | Cell-ID |

**Additional file: Table**. Mass cytometry panel for analysis of synovial cells.
