## Supplementary Materials for "High dimensional analyses of cells dissociated from cryopreserved synovial tissue"

**Additional File: Suppl Figure**
**Donlin, Rao, et al.**

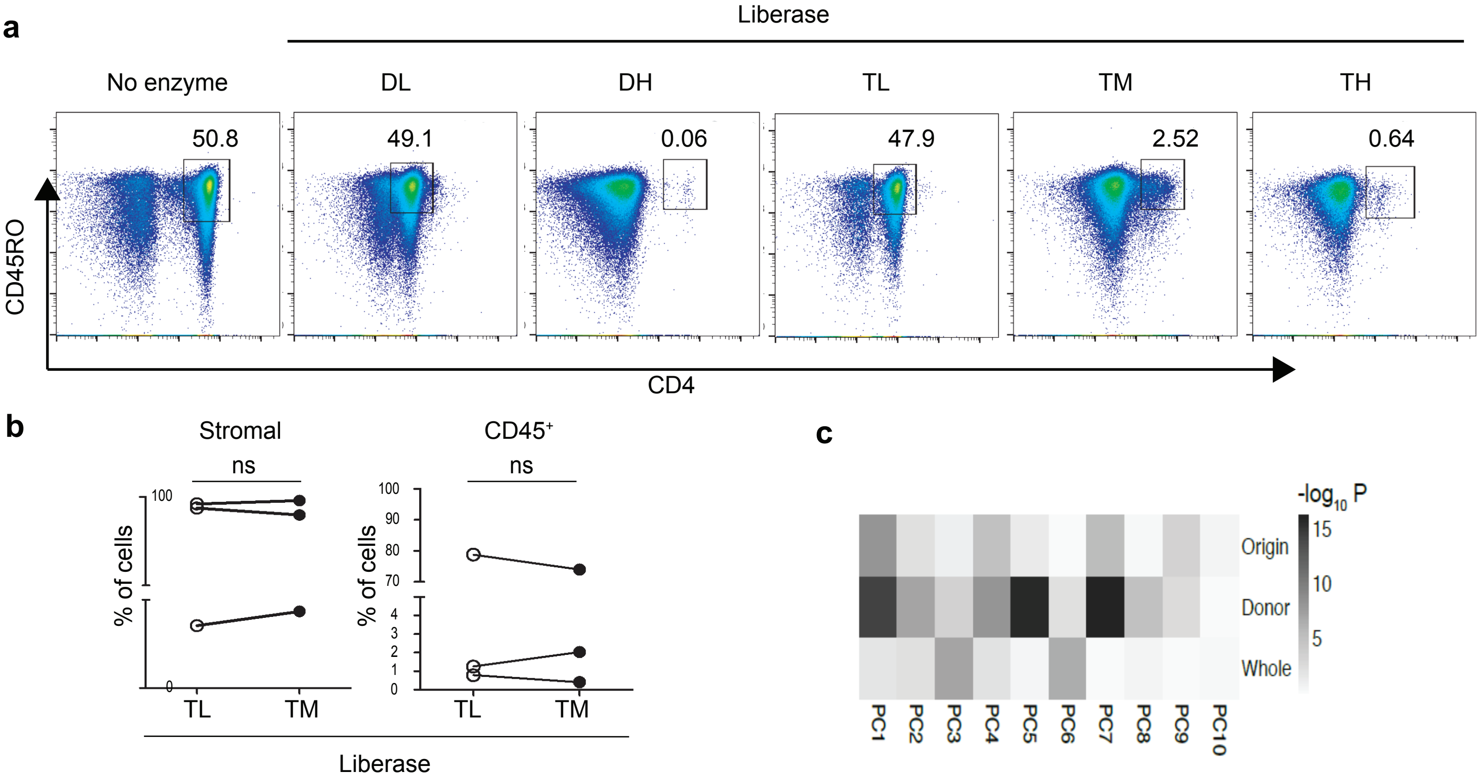


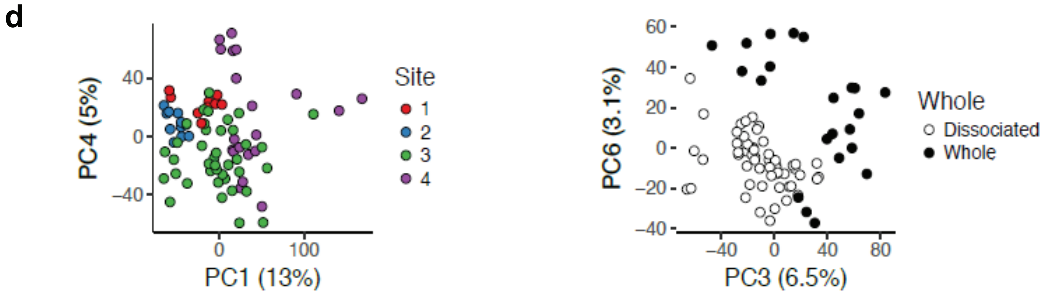


e

| Gene | Fold | Adj p-value |
|---|---|---|
| *DNAJB1* | 2.1 | 4.78E-27 |
| *ATF3* | 3.1 | 8.07E-25 |
| *JUN* | 2.3 | 9.31E-24 |
| *HSPA6* | 2.3 | 3.56E-23 |
| *HSPA1B* | 2.3 | 3.56E-23 |
| *FOSB* | 2.5 | 3.67E-18 |
| *HSPA1A* | 1.9 | 5.54E-18 |
| *FOS* | 2 | 7.16E-12 |
| *ZFP36* | 1.4 | 1.04E-09 |
| *EGR1* | 1.8 | 1.36E-08 |

**Additional file: Supplemental Figure. Additional variables in synovial tissue enzymatic treatment.** a) Flow cytometry analysis of human peripheral blood mononuclear cells after incubation with Liberase proteolytic enzyme formulations. Cells were first gated for viability and CD3 T cell receptor expression. b) Synovial tissue collected during arthroplasty surgery was mechanically disrupted and treated with or without a panel of Liberase proteolytic enzymes. Dissociated cells were analyzed by flow cytometry. Representative data from four biological replicates. ns, not significantly different. c) Principal component scores for variables in synovial tissue processing bulk (nonsorted) synoviocyte RNA-seq transcriptomics. d) Principal component analysis on disaggregated synoviocyte and whole tissue transcriptomics, color coded based on the clinical collection site for each sample (left panel) or whether the sequencing was from dissociated or whole/intact tissue. e) Stress response genes expressed higher in dissagregated synovial samples compared to whole tissue.The heatmap plots the –log10 P values associated with each factor for each principal component.
